# Signal Peptide Efficiency: from High-throughput Data to Prediction and Explanation

**DOI:** 10.1101/2022.05.13.489517

**Authors:** Stefano Grasso, Valentina Dabene, Margriet M.W.B. Hendriks, Priscilla Zwartjens, René Pellaux, Martin Held, Sven Panke, Jan Maarten van Dijl, Andreas Meyer, Tjeerd van Rij

## Abstract

The passage of proteins across biological membranes via the general secretory (Sec) pathway is a universally conserved process with critical functions in cell physiology and important industrial applications. Proteins are directed into the Sec pathway by a signal peptide at their N-terminus. Estimating the impact of physicochemical signal peptide features on protein secretion levels has not been achieved so far, partially due to the extreme sequence variability of signal peptides. To elucidate relevant features of the signal peptide sequence that influence secretion efficiency, an evaluation of ~12,000 different designed signal peptides was performed using a novel miniaturized high-throughput assay. The results were used to train a machine learning model, and a post-hoc explanation of the model is provided. By describing each signal peptide with a selection of 156 physicochemical features, it is now possible to both quantify feature importance and predict the protein secretion levels directed by each signal peptide. Our analyses allow the detection and explanation of the relevant signal peptide features influencing the efficiency of protein secretion, generating a versatile tool for the in silico evaluation of signal peptides.

## INTRODUCTION

The general protein secretion (Sec) machinery and its associated signal peptides (SPs) are responsible for the translocation of the majority of bacterial, archaeal and eukaryotic proteins across the cytoplasmic or endoplasmic reticular membranes (1–4). Because of its high capacity, the Sec pathway of various microorganisms was engineered to generate cell factories for the production of secreted proteins (3, 5, 6). Monoderm Gram-positive bacteria, like *Bacillus subtilis*, are preferred for this purpose, as products only need to pass a single membrane, which eases secretion and subsequent recovery of bulk amounts of protein from the fermentation broth (2, 7, 8).

The SP at the N-terminus targets a protein of interest (POI) to the extracellular space. It is composed of a positively charged N-region, a hydrophobic *α*-helical H-region, and a C-region that encompasses an Ala-X-Ala motif (with X being any amino acid) for cleavage by a signal peptidase during or after translocation (9–11). Thus, the presence of a SP within a protein sequence can be reliably detected (12). However, there are no tools to predict the secretion efficiency of a given SP-protein combination (13), in fact finding an efficient SP to secrete a POI is based on trial-and-error. Previous studies tested limited numbers of natural SP variants (i.e. up to 10^2^) and analyzed the relationships between secretion efficiency and some SP features (14–16). Such studies showed that the SP-POI match plays a crucial role in determining secretion efficiency (11), but did not unveil the underlying fundamental parameters.

Here we aimed to elucidate the relevant physicochemical features that determine secretion efficiency of a specific POI, namely the *α*-amylase AmyQ from *Bacillus amyloliquefaciens,*within a defined setting (i.e. specific growth and assay conditions), taking into account not only the amino acid (AA) but also the nucleotide sequences. To achieve our aim, we devised a workflow (Figure 1), based on the Design-Build-Test-Learn (DBTL) cycle approach. An SP-library was designed using 134 known wild-type SPs from *B. subtilis* as template. High-throughput (HT) quantification of secretion efficiency (17) was then used to generate a training dataset for a machine learning (ML) model. Finally, the impact of physicochemical SP features on the secretion efficiency was estimated using TreeSHAP (18–21) (hereafter SHAP).

**Figure 1.**
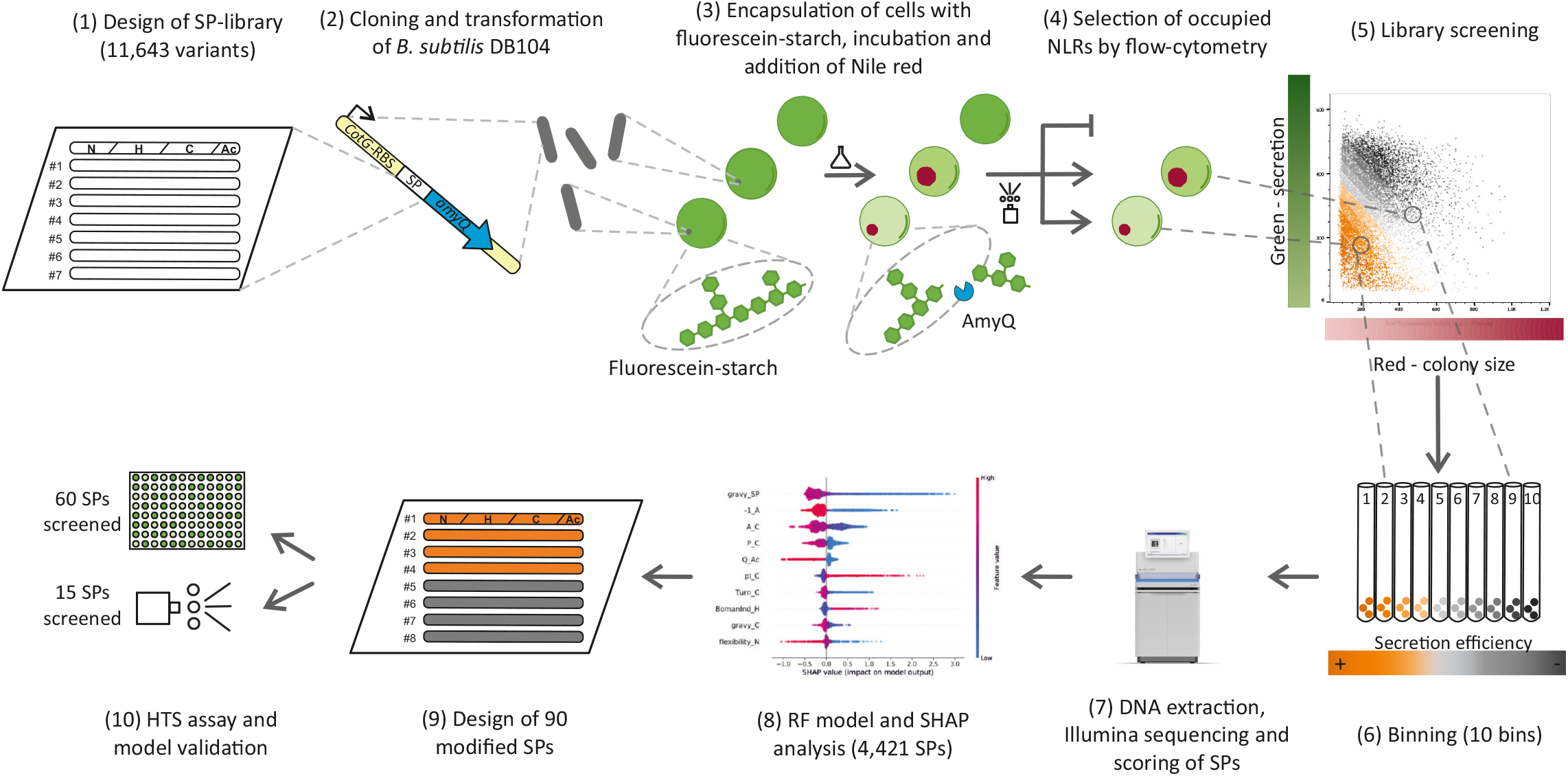
High throughput characterization of the signal peptide (SP) library. Experimental workflow: (1) A library of approximately 12,000 signal peptide (SP) variants was designed by modifying key features (e.g. charge, length, hydrophobicity); (2) the corresponding pool of oligonucleotides was cloned in frame with the sequence coding for mature AmyQ, and integrated into the *amyE* locus of *B. subtilis* DB104. (3) The clones were embedded in hydrogel beads, referred to as nanoliter reactors (NLRs), containing fluorescein-starch (mean diameter of 500 μm; average occupation of 0.3 bacterial cells per NLR). During incubation in culture medium, single cells grew into microcolonies and secreted AmyQ, which degrades the fluorescein-starch into (still fluorescent) low molecular weight fragments that are lost from the NLR by diffusion. After incubation, biomass in the NLRs was labelled by adding Nile red, a membrane-specific red fluorescent dye, and the NLRs were evaluated in 2 steps using a large particle flow cytometer. (4) Firstly, all empty NLRs were identified and discarded; (5–6) secondly, occupied NLRs were sequentially sorted into 10 bins, based on their green to red signal ratio. The green fluorescence signal is inversely proportional to the amount of secreted amylase (AmyQ) in the NLR; the red signal is instead directly proportional to the colony size. Therefore, clones with a high secretion efficiency are located in the lower left corner of the dot plot (5) and have a low bin number. (7) DNA from the NLRs of each bin was recovered and SP occurrence in any given bin was determined by NGS, leading to the construction of a frequency table of SPs across bins, used to calculate the secretion efficiency of each SP variant as a weighted average (WA). (8) WA values were subsequently combined with the features describing each SP to train a Random Forest (RF) regressor model. The RF model was then studied using SHAP for explanation and quantification of the impact of each feature on the model output (i.e. WA). (9) Information obtained by combining the RF model with the SHAP analysis was used to generate new SP variants with defined secretion levels to validate the model. (10) The designed validation sequences were processed following the same high-throughput (HT) screen, yet individually and not as a library. The secretion of amylase was quantified both with a microtiter plate assay (60 SPs) and by the NLR-based screening protocol (15 SPs), and the results were compared to the predictions.

## MATERIAL AND METHODS

### Library design

To identify the most relevant physicochemical features influencing the secretion efficiency directed by signal peptides (SPs), we designed a SP mutant library starting from 134 sequences (Supplementary Table 1) of known or highly probable *Bacillus subtilis* wild-type SPs. These SPs were initially selected based on literature (14) and via predictions by various computational tools (SignalP4.1 (22), SignalP3.0 (23), Phobius (24)). Next, the selected SPs were manually curated to remove false positives, knowing the final localization of their cognate native protein. As a point of novelty, we considered the SP sequences both as a single sequence (i.e. the whole SP) and as the juxtaposition of 4 separate parts, namely: the canonical N-, H-, and C-regions, plus a region referred to as ‘after cleavage’ (Ac-region), which consists of the first 3 amino acid residues (AAs) after the expected signal peptidase cleavage site. The Phobius tool for SP predictions was used to determine the boundaries of the 4 regions constituting each SP, still with partial manual curation based on evidence from literature. After defining the 4 regions for each SP, physicochemical properties were calculated for each region independently as well as for the complete SP. The 227 calculated properties are listed in Supplementary Table 2 while the respective methods of calculation and further explanations are reported in Supplementary Table 1.

From each of these 134 SPs, 94 mutant sub-libraries of 134 elements each were created. In each sub-library only one feature at a time was edited, while modifications to other interdependent features (e.g. the charge of an AA sequence affects also its isoelectric point and hydrophobicity) were minimized. Edited features at the AA level were hydrophobicity, charge, and length; edited features at the nucleotide level were codon usage and RNA secondary structures. The full list of varied features is presented in Supplementary Table 2. For each selected feature, multiple target levels (usually 4 or 5) were chosen. The rationale for selecting target levels was to allow for some expansion of the investigated design space without diverging too much from the biologically meaningful space of the wild-type SPs. The resulting SP-library was composed of the 94 sub-libraries and included a total of 11,643 unique sequences, which are presented in Supplementary Table 1.

### pSG01 plasmid construction

The plasmid pSG01 (Supplementary Figure 1, and see Supplementary Table 3 for the full plasmids list) was developed within this study in order to be used as a chromosomal integration vector for expression of the SP-library. To this end, the previously constructed genome-integrating vector pCS75 (25) (Supplementary Table 3) was cleaved with PmeI and EagI (NEB), the resulting fragments were separated on a 0.8% agarose gel, and the 7.8 kbp band, delimited by two regions homologous to the *B. subtilis amyE* gene, was excised and purified with the QIAquick Gel Extraction kit (Qiagen). The DNA sequence encoding the AmyQ mature protein (P00692) (i.e. without its SP) was ordered as a single gBlock G1 (Integrated DNA Technologies, Inc.) (see Supplementary Table 3 for full nucleotide sequence), amplified with primers P1 and P2 (see Supplementary Table 3 for a full primer list), digested with the same restriction enzymes as the vector, and purified with the DNA Clean & Concentrator-25 kit (ZymoSearch). The two DNA fragments were ligated, and the ligation mix was directly used to transform 10-beta competent *Escherichia coli* cells (NEB), to amplify pSG01. The resulting plasmid was verified and used to transform *dam*^-^/*dcm*^-^ competent *E. coli* cells (NEB), from which demethylated pSG01 was obtained for all downstream applications to increase the efficiency of *B. subtilis* transformation (26). Notably, 5’ to the SP-less *amyQ* gene, plasmid pSG01 contains two BsmBI (a type IIS restriction enzyme) restriction sites at 11 nt distance, which are oppositely oriented so that cleavage occurs upstream of each restriction enzyme recognition sequences, thus allowing for scar-less insertion of properly oriented DNA fragments. Moreover, this feature allows for the insertion of multiple DNA fragments in one step. After transformation of *B. subtilis*, pSG01 will integrate into the *amyE* gene, thereby disrupting the main source of amylase activity in *B. subtilis*.

### Expression strains and cloning of the library

*B. subtilis* strain DB104 (27), which lacks two major extracellular proteases, was selected to produce the library of designed SPs fused to AmyQ.

To obtain the final SP-library, pSG01 was endowed with two DNA fragments, using the two BsmBI restriction sites upstream of the SP-less *amyQ* gene: one fragment contained the *P_veg_* promoter (28), the native mRNA stabilizer of *cotG* (29), and a strong RBS from the *pre(mob)* gene of pUB110 (30), obtained as a single gBlock G2 (Integrated DNA Technologies, Inc.; Supplementary Table 3); the other fragment coded for one of the 11,643 designed SPs (ordered as an oligo pool from Twist Bioscience). Both fragments were designed to be amplified with P1 and P2 primers (Supplementary Table 3) and to present two terminal BsmBI cleavage sites generating complementary sticky ends to the vector for sequential assembly. Cloning was carried out using the StarGate (31) methodology and the resulting construct constitutively expressed the gene coding for the mature AmyQ fused at the N-terminus with one of the 11,643 designed SPs. A total of 3 mL StarGate reaction was mixed with 63 mL of competent *B. subtilis* DB104 that has been prepared using a modified Spizizen protocol (25). After 1 h of recovery at 37°C and 250 rpm, cells were plated on 62 Q-trays (Nunc™ Square BioAssay Dishes product n. 240835, ThermoFisher) each containing 200 mL of 2xPY medium (16 g/L peptone, 10 g/L yeast extract, 5 g/L NaCl) supplemented with 15 g/L agar and 300 μg/mL spectinomycin. After cell plating, the Q-trays were incubated at 30°C for 20 h.

The total number of grown colonies was estimated using a QPix 450 (Molecular Devices) automated microbial screening system. Two rounds of transformation were performed in order to obtain approximately 160,000 colonies, corresponding to a 10X coverage of the SP-library, and estimating 10% of clones containing pSG01 without inserts (data not shown). Plates were scraped to collect all colonies and rinsed with 2xPY. The collected cells were then transferred to several 50 mL Falcon tubes, mixed, and concentrated by centrifugation at 3,000xg for 5 min. The pellets were resuspended in 2xPY, the cell suspensions were pooled, thoroughly mixed, and supplemented with glycerol to a final concentration of 10% (v/v). The glycerol stock was aliquoted, snap frozen, and stored at −80°C. The cell concentration in the glycerol stocks, as determined by the optical density at 600 nm, was approximately 5.8*10^9^ cells/mL.

### Substrate preparation for NLR-based amylase assay

Dry corn starch (Sigma Aldrich, S9679) was re-suspended in 90/10 DMSO/water (v/v) to a final concentration of 2% (w/v), boiled for 30 min and allowed to cool to room temperature. An aliquot of 100 mL of the prepared solution was basified with 1 M NaOH until it reached a pH ≥ 9, then mixed with 1 mL of the reactive dye 5-([4,6-dichlorotriazin-2-yl]amino)fluorescein hydrochloride (DTAF) (Sigma Aldrich), previously dissolved in DMSO (20 mg/mL). After 1 h incubation at room temperature, the solution was neutralized with glacial acetic acid to stop the reaction, and the fluorescein-starch was precipitated with ethanol to remove the remaining free dye. The precipitated starch was resuspended in DMSO and subsequently ground with glass beads at 30 Hz for 20 min (Retsch). The resulting fluorescein-starch preparation was stored at 4°C and used as the substrate employed to monitor amylase activity within the nanoliter reactor (NLR)-based assay described below (32, 33).

### Cultivation of strains in NLRs

NLRs were synthesized starting from a mix of bacterial glycerol stocks, fluorescein-starch and sodium alginate, which was processed through a laminar jet break-up encapsulator (Nisco Engineering) to generate a monodisperse bead population. To prepare the mix, 200 μL of fluorescein-starch (4% w/v in DMSO) were diluted in 2 mL of resuspension medium (4 g/L yeast extract, 1 g/L tryptone, 20 mM TRIS pH 7) and added to 16 mL of sodium alginate 2.5% (w/v) aqueous solution. The number of bacterial cells to be included was defined to achieve an average occupation of 0.3 cells per NLR. To this end, the corresponding volume of the bacterial glycerol stock was added to the resuspension medium to reach a final volume of 2 mL, which was then mixed with the fluorescein-starch alginate preparation.

For NLR formation, the encapsulator was equipped with a 150 μm nozzle, and operated with a flow rate of 3.3 mL/min and a frequency of 650 Hz (34). This delivered NLRs with an average diameter of 500 μm (corresponding to a volume of approximately 65 nL). NLRs were allowed to harden for 15 min in 100 mM aqueous CaCl_2_, then isolated using a cell strainer (100 μm mesh size, Falcon, Becton Dickinson) and washed once with 10 mM aqueous CaCl_2_. NLRs were transferred into growth medium (4 g/L yeast extract, 1 g/L tryptone, 20 mM TRIS pH 7, 4 mM CaCl_2_, 300 μg/mL spectinomycin) with 0.5% (v/v) amylopectin to a final concentration of 100 g wet NLRs/L in Erlenmeyer flasks. The reactors were incubated in a shaker (150 rpm, room temperature) for approximately 13 h to allow cells to grow into microcolonies. NLRs were then recovered and washed twice with screening buffer (10 mM CaCl_2_, 10 mM TRIS pH 8). During each wash, the beads were allowed to sediment in a 50 mL Falcon tube, the supernatant discarded, and buffer added to achieve a concentration of 12.5 g of wet NLRs/L. Prior to screening, 40 μL of Nile red (Chemodex) (1 g/L in 90/10 DMSO/water, v/v) were added for every gram of wet NLRs to fluorescently stain the cells. The NLRs were incubated for 20 min under gentle shaking, washed once more with the screening buffer to remove surplus dye, and then subjected to flow cytometry and microscopic analysis. Bright-field and fluorescence microscopy images were recorded using an Axio Observer II with an AxioCam MR3 camera (Carl Zeiss Microscopy) to control for proper NLR synthesis and cell growth. For a detailed description of the flow cytometry analysis, see the section below.

If alginate beads with known SPs variants needed to be incubated together in the same vessel (to guarantee identical incubation conditions) and differentiated later in the flow cytometry analysis, the NLRs were synthesized with different concentrations of Pacific-blue (Ex 410 nm, Em 455 nm) labelled amino dextran (AD). Two concentrations, corresponding to 12 and 2.4 μL of the Pacific-blue AD stock solution (20 mg/mL in 0.2 M sodium bicarbonate, pH 8.3) per mL of fluorescein-starch alginate, were added. This polymer is not a substrate for AmyQ (data not shown) and does not interfere with the recording of fluorescein-based fluorescence (Ex 492 nm, Em 516 nm). Instead, the Pacific-blue content can be read out in the violet spectrum. Conjugation of the dye to the polymer was achieved by adding 5 mg of the amine-reactive Pacific Blue succinimidyl ester (ThermoFisher) to a solution of 20 mg AD (Fina Biosolutions) per mL of 0.2 M sodium bicarbonate (pH 8.3). The reaction was incubated for 6 h at room temperature. Then TRIS pH 7 was added to a final concentration of 50 mM to stop the reaction, and the solution was aliquoted and frozen.

Throughout the study, different *B. subtilis* strains, all generated in the same fashion and with the same vector, were analyzed using the NLR-based amylase assay. These included: 1) *B. subtilis* producing AmyQ with its native signal peptide (positive control, PC), 2) *B. subtilis* carrying the empty vector, without an inserted signal peptide (negative control, NC), 3) *B. subtilis* transformed with the SP-library, fused to AmyQ, or 4) *B. subtilis* producing AmyQ with SP variants with defined modification. The PC (1) and the NC (2) strains, and two variants producing AmyQ with known SPs were used to estimate the dynamic range and sensitivity of the NLR-based amylase activity assay (Figure 2a and Supplementary Figure 2). 15 strains producing AmyQ with SP variants with defined modification (4) were encapsulated and used to validate both the NLR-based screening assay and the model.

**Figure 2.**
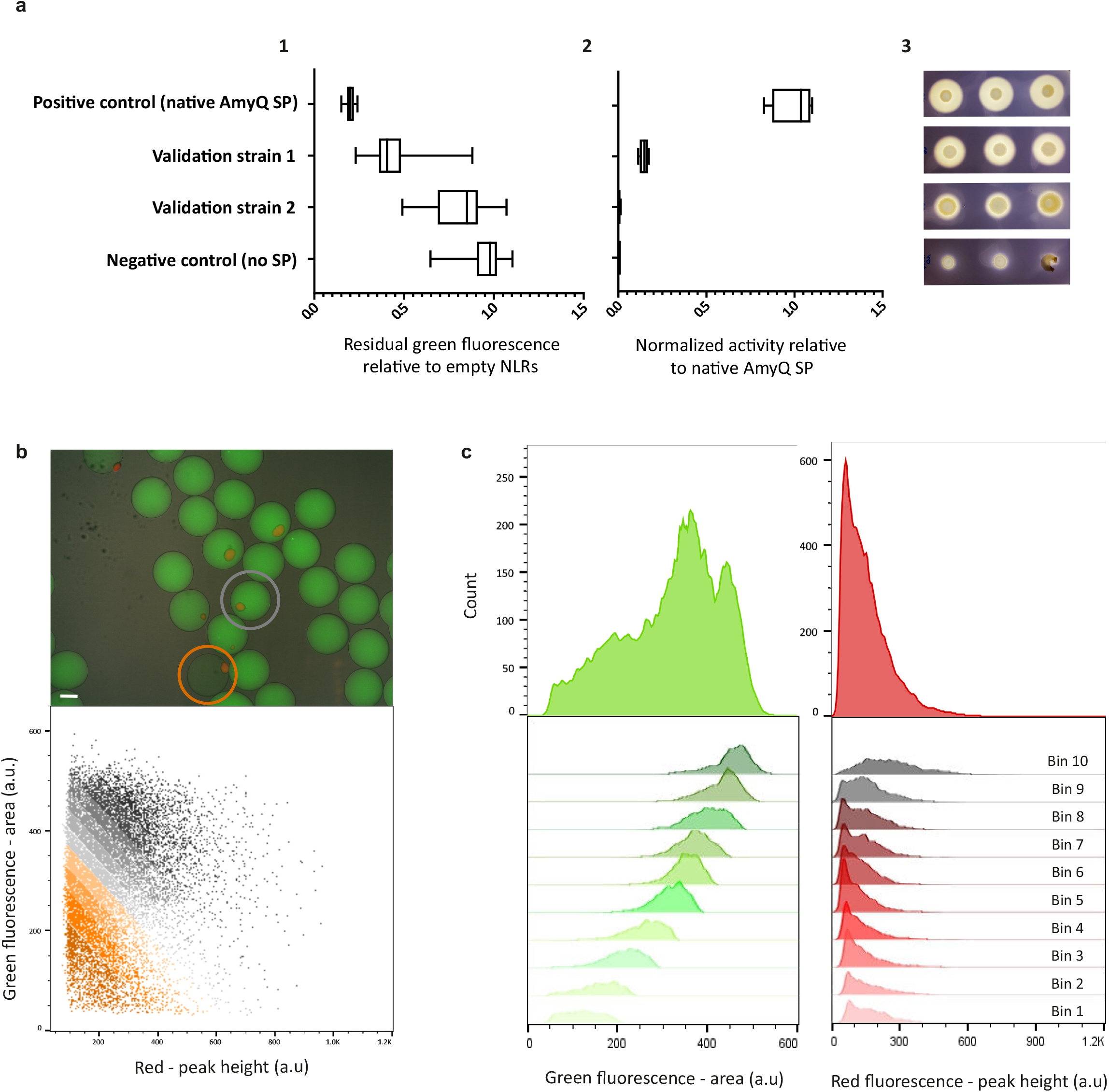
High-throughput of SP library screening in nanoliter reactors. (a) For initial validation of the NLR-based α-amylase assay, four *B. subtilis* strains secreting AmyQ to different levels were used: three strains with known SP amino acid sequences at the N-terminus (one of them with the native SP of AmyQ; positive control) and a *B. subtilis* strain synthesizing the amylase without an N-terminal SP (negative control). Amylase secretion of each strain was assessed using (1) the NLR-based assay, (2) the MTP colorimetric assay, and (3) the starch hydrolysis test. (1) For the NLR-based assay, the values represent the residual fluorescein-labelled starch still present in the occupied NLRs after cell growth, relative to the green fluorescence of the empty NLRs in the same population (set as 1). The recorded events were: positive control, 24 occupied and 850 empty NLRs; validation strain 1, 99 and 4329; validation strain 2, 50 and 932; negative control, 124 and 859. (2) For the MTP assay, the values are calculated relative to the amylase activity produced by the positive control (having a value of 1) and four biological replicates were performed. (3) The starch hydrolysis tests based on the starch-iodine reaction^23^. (b) Top: Overlay of bright-field and fluorescence microscopy images of NLRs after incubation in medium. Empty NLRs (no red dot) show a homogenous green fluorescence profile (no starch degradation), while NLRs harboring a colony (red dot) show different degrees of fluorescein-labelled starch degradation (orange circle: high secretor; grey circle: low secretor). Scale bar: 200 μm. Bottom: Dot plot representing all occupied NLRs from one experiment (approximately 20,000 NLRs). The gating applied during the second sorting step is depicted in orange-grey colour codes, which defines bins with distinct AmyQ secretion levels. (c) Green and red fluorescence profiles of all sorted events from the same experiment, both as a whole population (i.e. occupied NLRs; top panel) and divided into the 10 equally-sized bins (lower panel).

### NLR-based screening

The NLR-based screening of the clones carrying the SP-library, was performed with a large particle flow cytometer, which allowed to read out the amount of starch, of cells, and, if applicable, of amino dextran in each NLR, based on different fluorescence signals. Specifically, we used a BioSorter (Union Biometrica) to record for each NLR green (excitation laser 488 nm, beam splitter DM 562, emission filter BP 510/23 nm), red (excitation laser 561 nm, TR mirror, emission filter BP 615/24 nm) and violet fluorescence (excitation laser 405 nm, beam splitter DM 495, emission filter BP 445/40 nm).

Each screening round was performed in two sequential steps. During the first step, all events were analyzed in bulk mode, at a maximum of 90 Hz, and NLRs with a positive red fluorescence (peak height, i.e. presence of colonies stained with Nile red) were sorted into a 50 mL Falcon tube, containing 5 mL of screening buffer. The isolated population represented approximately 20% of all the NLRs, in agreement with the occupation estimated from the cell concentration in the glycerol stocks. Prior to the second step, the values of green and red fluorescence of each sorted NLR were graphically visualized using the FlowPilot software provided by the BioSorter manufacturer. The graph was then used to divide all events in 10 bins based on the ratio between green fluorescence (area, representative of amylase activity and secretion levels) and red fluorescence (peak height, representative of total biomass). The bin width was thus adjusted to have 10% of the events sorted in step 1, in each bin. For the second step, sorted NLRs were run through the Biosorter ten consecutive times, every time isolating in bulk mode the events falling in one bin. In particular, the sorting started from the bin with the lowest green to red ratio (i.e. highest secretion/biomass ratio), bin 1, and then moved progressively to bins with higher green/red ratios. The screening analysis was repeated until the number of occupied NLRs (i.e. positive red fluorescence) reached 160,000, to ensure a 10x coverage of the SP-library. Additionally, after cell encapsulation and growth in the NLRs, 53,588 occupied NLRs were sorted in 3 rounds, without performing any further binning, and treated separately. This sample, named hereafter ‘occupation control’, was processed and sequenced with the 10 bins, and later used to gather information about the library coverage and the *B. subtilis* population at this step of the workflow.

To recover the NLR-embedded cells, binned samples were incubated for 10 min under gentle shaking with 2xTY medium (16 g/L tryptone, 10 g/L yeast extract, 5 g/L NaCl) supplemented with potassium phosphate buffer (pH 7) to a final concentration of 0.2 M, at which point full dissolution of the cross-linked calcium alginate had been achieved. Bacterial cells were pelleted by centrifugation (4,000xg, 30 min), the supernatant was discarded, and the pellet stored at −80°C.

### Genomic DNA extraction and NGS library preparation

Samples from the 10 bins, the occupation control, and the initial glycerol stock were thawed on ice, centrifuged for 1 min at 16,000xg, and the supernatant was discarded. Afterwards cells were resuspended in 0.85% (w/v) aqueous NaCl, supplemented with 250 μg/mL of RNase A (Macherey-Nagel) and 0.5 mg/mL of lysing enzymes from *Trichoderma harzianum* (Sigma-Aldrich, L4142). After incubating for 10 min at 37°C, EDTA and SDS were added to final concentrations of 15 mM and 1.2%, respectively. Samples were vortexed thoroughly, ammonium acetate was added to a concentration of 2.5 M, and then samples were vortexed again. Precipitated proteins were pelleted by centrifugation at 22,000xg for 15 min at 4°C. The supernatant was transferred to a fresh reaction cup, supplemented with an equal volume of 2-propanol and gently mixed. DNA was then pelleted by centrifugation at 22,000xg for 40 min at 4°C. The supernatant was discarded and the pellet was washed twice with ice-cold 70% ethanol, dried, and resuspended in 10 mM Tris-HCl pH 7.5.

Each sample was then amplified by PCR, using Phusion polymerase (NEB), with primers P3-P15 (Supplementary Table 3) that anneal immediately up- and downstream of the inserted SP sequence in pSG01, adding barcodes to identify the sample (primers P3-P15 in Supplementary Table 3, containing Illumina Nextera tagmentation adapters and, in each forward primer, a specific barcode). PCR products were then purified and recovered in milliQ water. Amplicons were analyzed with a bioanalyzer (LabChip GXII, Caliper Life Sciences) using a 5K HT DNA chip, to check size and concentration of the fragment. The 12 PCRs products, corresponding to the 10 bins and the two controls, were pooled and sequenced as a Nextera library (Illumina) by the company BaseClear B.V. (Leiden, NL) on a NovaSeq machine (Illumina) in paired ends, for a total of 26,175,197 2×150bp reads. For both forward and reverse raw reads, the Phred scores had an average of 36 and a median of 37.

### Reads pre-processing and mapping

The software FastQC version 0.11.8 (35) was used for quality inspection of the sequencing data. First, possible adapters were removed from the 3’ end of the reads (read-trough adapters), since they could confound the merging process when the read length and insert size are comparable. To this end, the software NGmerge (36) version 0.2dev was used in “adapter removal” mode, with 0 mismatches allowed. Sequences were thus merged into longer pseudoreads using PEAR (37) version 0.9.11, with a minimum overlap of 5nt and a p-value of 0.001. This yielded 26,105,901 pairs of reads (99.735% of the total reads) to be merged, with the remaining reads unassembled and no read discarded. Pseudoreads were then sorted in the 10 bins and the two controls, based on the respective barcodes, using the ‘fastx_barcode_splitter.pl’ script from FASTX-Toolkit (38) looking only at the 5’ (‘*--bol*’ option) and allowing only 1 mismatch. This resulted in 25,980,025 (99.254% of the total reads) demultiplexed pseudoreads. Any remaining adapter (including the barcode) at both 5’ and 3’ of the assembled reads was removed using cutadapt (39) version 2.3, without any read loss. The obtained pseudoreads were then mapped to the reference sequences (i.e. the designed SPs) using BBMap (40) version 37.93, with *‘perfectmode’* activated; and behavior for ambiguously mapped reads was set to ‘best alignment’ (Supplementary Table 4). Occurrences for each bin and both controls were counted for each of the designed sequences and the resulting frequency table was later used for model construction (Supplementary Table 1).

### Data preprocessing, feature extraction, model construction and interpretation

To identify the possible influence of investigated features on protein secretion, we decided to train a simple machine learning model. This procedure, combined with an interpretation of the model, would allow us to obtain a predictive model that could yield important mechanistic insights into the features determining the secretion efficiency of different SPs.

First of all, sequences with low abundance, corresponding to less than 255 reads in the most populated bin, were discarded. This resulted in 4,421 informative SPs, which were used to train and test the model. As a different number of NLRs was collected for each bin, the occurrence of reads was normalized across bins so that they contained the same number of NLRs. To score SPs, we assumed that bins were equidistant and each bin had an average value corresponding to its number. A weighted average (WA), i.e. the summation of bin values weighted on the relative frequencies of reads, was calculated for each SP and used as a secretion score. The WA values of selected SPs could thus range from 1 (i.e. the best secreting SPs with all occurrences detected in bin 1) to 10 (i.e. the worst secreting SPs with all occurrences detected in bin 10).

From the 227 calculated features, 22 were discarded because they showed no variation either in the designed SP-library or in the informative SPs dataset, which was a subset of the designed library (Supplementary Table 2). Furthermore, it was decided to minimize the number of features with a correlation coefficient higher than 0.7 to avoid a spread of importance, as attributed by the model, among them. Thus, out of the initial 227 features, 96 were retained, while 110 features, with correlation coefficients higher than 0.7, were selected for clustering. Clustering was carried out through affinity propagation (41) using the scikit-learn (42) python package with standard parameters. Notably, affinity propagation was selected as the clustering algorithm, because of its intrinsic capability of inferring the total number of clusters. This resulted in 14 clusters of correlating features, out of which 22 features were selected and added back to the feature set. Specifically, for 12 of the clusters the centroid was selected, for 1 cluster the centroid and an additional feature were selected, and for the last cluster with lowly correlating features all of the 7 features were included. Altogether, this procedure resulted in a total of 116 features, to which 40 Boolean dummy variables were added that describe the AAs in positions −3 and −1, respective to the signal peptidase cleavage site. This resulted in the final selection of 156 features describing the selected SPs (Supplementary Table 2). In order to verify that the selected features were relevant (i.e. provided variance) within the datasets, and thus needed to train the model, a principal component analysis was performed on: (i) the 134 wild-type known SPs, (ii) the set of 11,643 unique SP (i.e. the SP-library), and (iii) the set of 4,421 informative SPs (Supplementary Figure 3).

To construct the model, the matrix of 4,421 informative SPs and 156 features was used as the independent variable, while the array of 4,421 WA values was used as the dependent variable. These matrices were split with the Kennard-Stone algorithm (43, 44) into a training set and a test set of 3,095 and 1,326 informative SPs, respectively. From available models, a Random Forest (45, 46) (RF) Regressor model from scikit-learn (42) was implemented. To identify the best hyperparameters a 5-time cross-validation grid search was performed using the training set. From the 15,435 tested combinations of hyperparameters, the following set of hyperparameters was selected, which balance the predictive power and the size of the model: max depth 25, max features 156, min samples leaf 0.0001 of the training set, min samples split 0.001 of the training set, and estimators 75. The model was subsequently evaluated on the test set, and scored calculating the mean squared error between measured and predicted values (Figure 3a).

**Figure 3.**
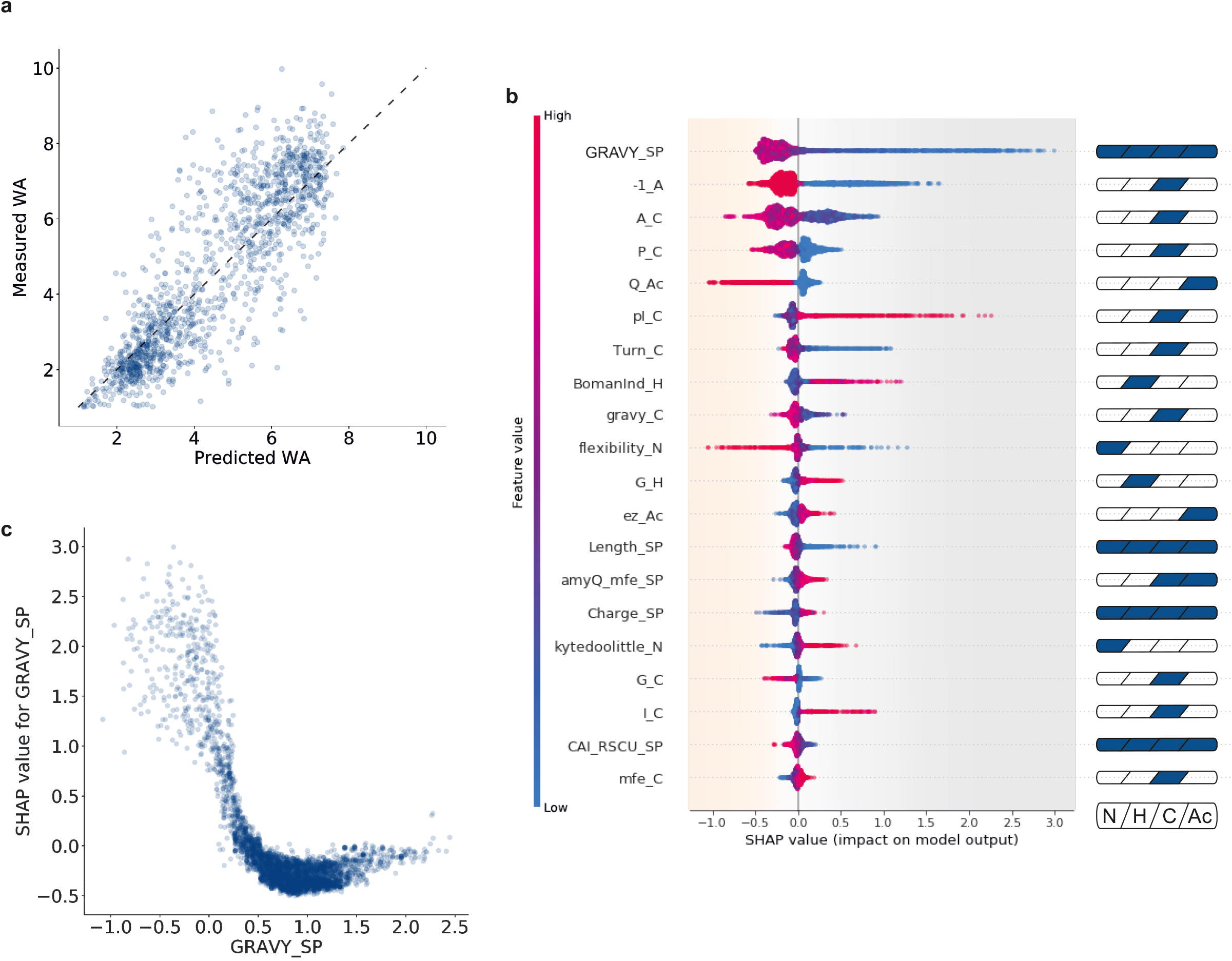
Model explanation. (a) Predictions of the trained Random Forest (RF) regressor model on the test set. On the ordinate WA values for SPs belonging to the test set and measured with the NLR-based amylase assay are reported; on the abscissa their WA values predicted with the trained RF model are displayed. Note that measurements and prediction show a high degree of agreement with a calculated mean squared error of 1.22 WA, thus indicating a good performance of the generated RF model. The dashed line represents the ideal situation where all predicted and measured values would align. (b) SHAP summary plot of the 20 most impactful features, where a high dispersion of SHAP values on the abscissa indicates a broad effect of the respective feature on the model. Each data point represents a specific SP, the color of the data point indicates the value of that feature in the feature-specific scale, and the position on the abscissa indicates the SHAP value for that particular feature. SHAP values for the whole dataset sum up to the base value of the model (4.45 WA, average model output calculated over the 4,421 selected SPs). Positive SHAP values indicate a negative impact on the model outcome, and vice versa. A brief description of each feature is presented as a label on the right (for full descriptions see Supplementary Table 2). Cartoons on the right highlight the corresponding SP parts of each particular feature. (c) SHAP-dependence plot for ‘GRAVY_SP’, which is a two-dimensional representation of the information summarized by the first line of panel b. The GRAVY index is represented on the abscissa: negative values indicate low hydrophobicity, and positive ones high hydrophobicity. On the ordinate SHAP values are displayed: negative values indicate a beneficial effect on protein secretion, and positive ones indicate a detrimental effect. Vertical dispersion of SHAP values for similar GRAVY indexes can be explained through the interaction effect between features (described by SHAP interaction values, summarized in Supplementary Figure 8 and Supplementary Figure 9). To exemplify, the high variability visible in the negative range of the GRAVY index is to be attributed mainly to the feature ‘-1_A’ (Supplementary Figure 7). The data imply that a very low hydrophobicity will have a strong negative impact on protein secretion, while a GRAVY index value of around 1.0 will be most favorable for protein secretion.

The RF model was analyzed to gain mechanistic insights and an explanation of the model itself. For this task, the TreeSHAP method from the SHAP (SHapley Additive exPlanation) (18–20) package was used since, being based on Shapley values, it is advantageous in terms of consistency, allows for a more reliable comparison of feature attribution values, and allows users to understand the model explanation. A further advantage of SHAP is that it includes both local and global explanations, thereby providing explainability for both the whole dataset and the individual SPs. Nevertheless, it should be emphasized that SHAP only provides an explanation of the model based on the contribution of individual features to the final output. SHAP does not necessarily uncover the causal relationships between individual SP features and the actual protein secretion efficiency as displayed by a bacterial cell. Furthermore, is noteworthy that SHAP provides the possibility to determine the type of relationship between each individual feature and the predicted output, and to determine second order interactions that occur between features.

### Assay validation

To show both the reliability of the NLR-based amylase activity assay and to assess the correctness of the model, we used two orthogonal procedures to measure the amylase activity from strains producing AmyQ with selected SPs: a commercial assay in 96-wells microtiter plates (MTPs) and a hydrolysis test on starch agar plates (47). For the MTP assay bacteria were precultured overnight into 2xPY supplemented with 300 μg/mL spectinomycin, subsequently diluted 100-fold in the same medium and grown for 7.5 h at 37°C and 250 rpm. Throughout the validation of the NLR-based amylase assay and of the model, two cultivation vessels were applied: deep-well plates filled with 300 μL of medium and/or Erlenmeyer shake flasks (25 mL culture volume in 250 mL flask). Aliquots of the different cultures were collected, the cells were pelleted, and 9 μL of the supernatants were mixed with 9 μL of Ceralpha reagent (Megazymes). The reaction mixtures were incubated for 20 min (standard version) or 90 min (sensitive version) at room temperature on a shaker (1,000 rpm) and then the reactions were stopped through the addition of 200 μL of 1% (w/v) 2-amino-2-(hydroxymethyl)propane-1,3-diol (Tris-base, pH 9). The amylase activity was then measured by monitoring the absorbance at 405 nm with a Tecan Infinite M200 Pro. Similar to the NLR-based assay, the optical density at 600 nm of the cultures was measured and used to normalize all samples for the biomass. Eventually, the OD-normalized absorption values were expressed relative to the amylase activity obtained from cultures that secreted AmyQ with its native SP (positive control, PC), defined as 100%, and to the amylase activity in the growth medium of a strain containing pSG01 without an inserted SP sequence (negative control, NC), defined as 0%.

For the hydrolysis test on starch agar plates, glycerol stocks of the selected variants were diluted 100-fold in 300 μL 2xPY supplemented with 300 μg/mL spectinomycin, and grown until they reached mid-exponential phase (6 h, 37°C, 250 rpm). An aliquot of 2 μL of the cell culture was spotted on 2xPY-agar plates supplemented with 300 μg/mL spectinomycin and 0.2% (w/v) potato starch (Sigma Aldrich). After overnight incubation at 37°C, the plates were flooded with Lugol’s iodine (Carl Roth), which interacts with starch and generates a dark color. Where starch is degraded, a clear zone arises, e.g. around a colony, and the area of this clear degradation zone is approximately proportional to the amount of amylase secreted (47).The standard MTP assay was used for initial validation of the NLR-based assay (Figure 2a, 2) and the screening strategy. To further validate the assay, 95 SP-AmyQ fusions, randomly picked from the 4,421 variants used to train and test the model, were subjected to the MTP assay (see activity values in Supplementary Table 5 and Supplementary Figure 4). As 72 randomly picked variants showed no amylase activity, presumably due to too low secretion levels, the sensitive version of the MTP assay was applied, and AmyQ activities could be determined for 15 more clones.

The starch hydrolysis on plate was also applied for the partial validation of the NLR-based assay (Figure 2a, 3) and if no amylase activity could be detected with the sensitive MTP assay (i.e. Abs_405_ < 0.1; Supplementary Figure 5).

### Model validation

To further validate our model, small sets of SPs were manually edited to tune their predicted secretion levels. 30 SPs directing high-level secretion of AmyQ (i.e. ‘good performers’) were manually edited until the model predicted them to direct AmyQ secretion with poor efficiency (Group 1). Similarly, another 30 SPs directing low-level secretion of AmyQ were edited in order to improve their efficiency (Group 2) (see Supplementary Table 6 for full list of SPs).

As an additional validation approach, we generated pseudo-random SP AA sequences, with a home-made script, described as follows. Based on 134 sequences of known and highly probable SPs, 7 dictionaries were calculated that map each AA to its relative frequency: one for the N-region (excluding the initial Met), one for the H-region, one for the C-region except the last 3 residues, one for each −3, −2 and −1 position relative to the signal peptidase cleavage site, and the last one for the Ac-region (i.e. positions +1, +2 and +3 together). Using these values, 10,000 sequences for each region were generated as follows: for the N-region a Met was always placed in front of a stretch of 1 to 10 residues built on the frequency table; the H-region simply consisted of a stretch of 9 to 16 residues built on the frequency table; the C-region was built juxtaposing a stretch of 4 to 11 residues to the 3 single residues for positions −3,-2, and −1, each based on its frequency table; and for the N-terminus of the mature protein after signal peptidase cleavage, a stretch of 3 residues based on its frequencies was built. To minimize the occurrence of SPs with features too far from the distribution of the known, or probably representing wildtype *B. subtilis* SPs, the Kolmogorov-Smirnov (48, 49) statistic test was applied, which compared the distribution of various features (data not shown). In case the number of similar distributions (considered as all those features with a calculated p-value above 0.1) was below a certain threshold, i.e. at least 21, 16, 18 and 17 features respectively for the N-, H-, C-, and Ac-regions, the batch of 10,000 sequences was discarded and the process was repeated. In such a way, the 4 designated regions (i.e. the N-, H-, C-, and Ac-regions) were built independently from each other and their possible combinations or interactions were not considered. When 10,000 sequences for each region were generated, they were juxtaposed to form a full SP. Then, sequences equal or longer than 33 AAs (calculating the length up to and including the Ac-region) were discarded, thus resulting in 4903 valid SPs. To generate the relative coding sequences, the AA sequences were retro-translated with an unambiguous dictionary where only the most frequent codon for each AA was present. Subsequently, having both a nucleotide and an AA sequence for each SP, the 156 features of the final model were calculated, and the respective SPs’ secretion efficiency (i.e. the WA value) was predicted using the RF model. A set of 32 SPs (Group 3) was selected with the requirement to be predicted by our model as very good secretors (see Supplementary Table 6 for full list of SPs). Eventually, the different features for each SP were also analyzed and explained through SHAP.

For manually edited as well as pseudo-randomly designed SPs, the respective SP-encoding sequences were ordered (Twist Biosciences), cloned in pSG01 and used to transform *B. subtilis* DB104, following the same procedure as applied for the generation of the SP-library. For the 61 successfully constructed SP-AmyQ fusions, the amylase activity was monitored in the MTP assay as described above. Additionally, 15 of these fusions (5 for each of the three groups) were further tested via the NLR-based amylase activity assay, to verify the model in the same experimental setup used to generate it (Figure 4).

**Figure 4.**
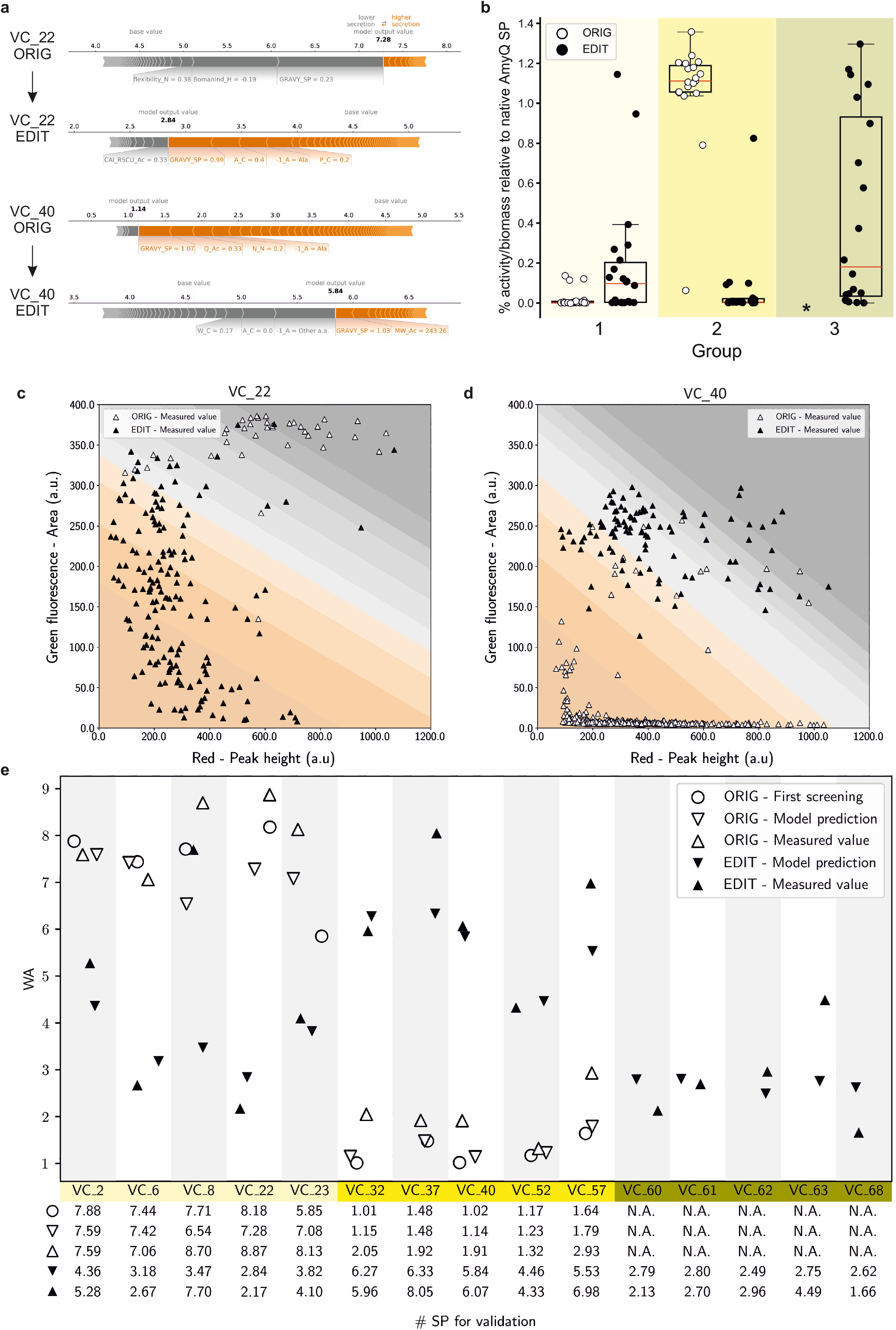
Assay and model validation. (a) SHAP force plots for SP variants VC_22 and VC_40 before (‘ORIG’) and after (‘EDIT’) editing. The impact of each of the most relevant features on secretion efficiency is quantitatively assessed. Each segment is sized proportionally to its impact on the model, their summation is equal to the difference between the base value (4.45 WA for all SPs) and the output value (i.e. the predicted WA value of each SP). Features colored in gray have a negative impact on the secretion efficiency of the specified SP, while features colored in orange have a positive impact. (b) Box plot showing amylase activity measured with the MTP assay for the 3 groups of SPs: Group 1, originally poorly secreting SPs (light yellow); Group 2, originally highly efficient SPs (yellow); and Group 3, pseudo-randomly designed SPs (dark yellow). The circles represent the 60 selected SPs before (white circles, “ORIG”) and after (black circles, “EDIT”) editing. The efficiency of the native SP of AmyQ is equal to 1 (* = not applicable, as the pseudo-randomly designed SPs were not present in the SP-library). (c) and (d) Dot plots showing the NLR-based analyses for *B. subtilis* strains secreting AmyQ with SPs VC_22 (Group 1) and VC_40 (Group 2), respectively. White triangles indicate NLRs harbouring strains secreting AmyQ with the original variant from the SP-library, while black triangles indicate NLRs with strains secreting AmyQ with the edited SP. In the background, the 10 different bins are indicated using the same colour code as in Figure 2b. (e) Summary plot of the model validation. For each of the 15 selected SPs, 5 data points are shown: open symbols indicate the original variant from the SP-library, while black symbols designate the engineered SP derivative. The open circles mark the WA measured during the initial screening, triangles pointing downward denote WA values predicted by the model, whereas triangles pointing upward denote WA values measured during model validation. Groups of SPs are highlighted in different shades of yellow as in (a). Below the graph, values are listed to allow a more detailed comparison.

## RESULTS AND DISCUSSION

### High-throughput SP library screening in nanoliter reactors (NLRs)

Starting from a selection of 134 known wild-type SPs from *B. subtilis*, a library of 11,643 unique SPs (Supplementary Table 1) was rationally designed to expand the sampling space and the variance of naturally occurring SP sequences. We individually modified 7 specific physicochemical features on 94 designated levels (Supplementary Table 2), while concomitantly minimizing their influence over related ones (e.g. editing the charge while avoiding a significant hydrophobicity change). Furthermore, each SP was treated both as a single sequence, and as 4 juxtaposed segments (i.e. the 3 aforementioned regions plus a short stretch of 3 AA after the cleavage site). The designed SP-library was then introduced into *B. subtilis* strain DB104 using a genome-integrating vector. A total of 160,000 clones was harvested, achieving a 10X coverage of the SP-library.

The secretion efficiency associated with each SP variant was determined by measuring amylolytic degradation of fluorescein-labelled starch upon encapsulation of the library strains in the so-called nanoliter reactors (NLRs) (32, 33). In this assay, green fluorescence of each NLR is rapidly measured via flow cytometry, allowing the assessment of secretion levels of active AmyQ (Figure 2 and Supplementary Figure 2). For initial validation, the high-throughput methodology was compared to two alternative assays: a microtiter plate (MTP) format using a synthetic substrate (Figure 2a, 2) and a starch hydrolysis test using agar plates (47) (Figure 2a, 3). The results of the different assays are comparable and show the highest sensitivity and dynamic range for the NLR-based assay.

Next, we performed the HT screening with the SP-library measuring simultaneously amylase activity and bacterial biomass. The latter was achieved by incubating the NLRs with Nile red, a hydrophobic red dye that interacts with cell membranes (50) and fluorescently labels the microcolonies. Occupied NLRs were separated in 10 equally populated bins, based on enzymatic activity per biomass unit (i.e. the ratio between green and red signals). Variant collection was followed by DNA sequencing to determine the abundance of each SP variant in each bin and, as a control, in the original library after transformation and before sorting the NLRs. Occurrence values were used to generate a weighted average (WA), assuming equidistance between bins, and this WA was used as an efficiency score for each SP.

As shown by sequencing, 92% of the 11,643 unique rationally designed SPs were successfully introduced into *B. subtilis*, while 83% were retrieved after screening (Supplementary Table 4). Such reduction may relate to SP-dependent impaired growth and the resulting high background-to-noise ratios for small colonies.

We subsequently characterized the sensitivity of the NLR-based secretion assay by comparing values from 95 randomly picked library clones in NLRs versus MTPs (Supplementary Figure 4). As 73 of the selected 95 variants could not be measured using the standard MTP format, we applied also a classical starch hydrolysis test on agar plates to verify the low-secreting variants (Supplementary Figure 5). These experiments highlighted the superior sensitivity of the NLR-based secretion assay.

### Machine Learning model to predict SP efficiency and model explanation

To test and train our ML model, we evaluated the number of physicochemical features in the SP dataset. Starting from an initial set of 267 features, 156 informative features were retained to describe each SP (Supplementary Table 2). This step removed features either presenting no variability or exhibiting a high correlation with another feature in the training dataset. A further reduction of dimensionality proved to be unnecessary, as the PCA analysis showed that each of the principal components contributed to the explained variation (Supplementary Figure 3). Additionally, the same number of components was necessary to describe the whole variance of the 11,643 unique rationally designed SPs and the 4,421 informative SPs, indicating that, despite the loss in the total number of data points, there was no loss in the variation of the dataset. In contrast, the PCA showed that, to explain the same variation, more principal components are needed within the designed library than for the wild-type set of SPs, underpinning the improvement of the assayed space gained with our design. The array of 156 features is thus to be considered as the independent variable, and the single value of secretion efficiency (WA), as the dependent one.

Three quarters of our dataset were used to train a random forest (RF) regression algorithm, resulting in a mean squared error (MSE) of 1.75 WA, while the remaining quarter was used as a test set, resulting in an MSE of 1.22 WA (Figure 3a). After this first validation, we proceeded to provide explanations for the RF model predictions. Due to the complexity in explaining such a developed RF model (51, 52), SHAP (18–20) was used to extract information about the importance of the features, and their interaction effects (Supplementary Figure 6).

Due to the large number of features fed into the model and the notable amount of information provided by SHAP, only the most relevant and representative findings are discussed. To fully explore the model, a Jupyter notebook and an interactive tool (File S1) are available as Supplementary Data. The 20 most impactful features in our model are shown in Figure 3b. Some of these features were already documented in literature (13, 53, 54), for instance the overall SP hydrophobicity (‘GRAVY_SP’), the helix-breaking residue at the end of the H-region (‘P_C’ and ‘Turn_C’), or the cleavage consensus sequence (e.g. ‘-1_A’ and ‘A_C’). Notably, even for such known features, the impact on secretion could so far only be qualitatively estimated based on their distributions in wild-type SPs. With our approach, a more precise quantification is now achieved, establishing favorable, neutral or detrimental values and their impact on the predicted secretion efficiency. Additionally, it is now possible to determine relationships between features and secretion efficiency (e.g. linear, sigmoidal, monotonic), as illustrated with the dependency plot for a simple feature, such as ‘GRAVY_SP’ (Figure 3c), whose wild-type distribution is known but only includes positive values (54, 55). Our model analysis shows that functional positive GRAVY values are favorable, while negative GRAVY values will be detrimental. Moreover, thanks to the applied segmentation approach (i.e. 4 juxtaposed regions for each SP), it is possible to visualize how particular features can have different relevance, depending on whether we consider the whole SP or only a single region. This is clearly exemplified by the feature ‘Charge_SP’, for which an overall value lower than +2 increases secretion efficiency, and a slightly negative charge is even more favorable. In contrast, inspection of the charge of the N-region, represented by ‘Flexibilty_N’ (Supplementary Table 2), shows that values close to +2 or higher favor secretion. Analogously, different features describing the same region can influence each other’s impact. For instance, the feature ‘BomanInd_H’, which positively correlates with charge and negatively with hydrophobicity (Supplementary Tables 1 and 2), shows that a high level of hydrophobicity (low ‘BomanInd_H’) in the H-region can favor secretion. At the same time, judged by the feature ‘G_H’ (Gly content in the H-region), it appears that high hydrophobicity due to a high Gly content is not favorable, most likely because Gly reduces the stability of *α*-helices. Because of the applied feature selection process, one feature (e.g. ‘BomanInd_H’) may be representative of similar ones (e.g. ‘pI_H’ and ‘Charge_H’), which sets a limit to our immediate understanding of the influence of some properties. Nonetheless, with the present approach, we can retain, explain and trace back to their correlating counterparts, physicochemical properties of SPs, rather than less biologically significant indicators (e.g. principal components of a PCA, or the ‘D-score’ from SignalP (14, 23)).

With SHAP it is possible to analyze and quantify pairwise interactions between features, which explains why equal values of the same feature can influence the model to different extents. For instance, the vertical dispersion of ‘GRAVY_SP’, which is especially pronounced for negative hydrophobicity values (Figure 3c), is to be attributed to the feature ‘-1A’ (Supplementary Figure 7). Furthermore, overall interactions seem to play minor roles in our model, as the most impactful interaction (‘Q_Ac’-‘A_C’) has limited impact on the overall output (Supplementary Figures 8 and 9). One possible explanation is that interactions occur at orders higher than the second, and as such would not be represented in the model.

### Assay and model validation through rationally edited and pseudo-randomly designed SPs

To validate our model, we decided (i) to rationally tune the secretion efficiency of screened SPs, and (ii) to *in silico* screen a library of pseudo-randomly designed SPs for high secretion levels. From the previously screened SPs we selected 30 sequences that poorly (Group 1) and 30 that highly (Group 2) directed AmyQ secretion, and we manually modified their nucleotide and amino acidic sequences to invert their efficiency (Figure 4a and Supplementary Table 6). An interactive exploration of original and edited SPs is possible through the File S2 (original) and File S3 (edited). Additionally, we generated 4,903 pseudo-randomly designed SPs, predicted their secretion efficiency, and picked 32 amongst the potentially best-performing SPs (average WA of selected SPs is 2.64) to be tested (Group 3). Out of the 92 SPs selected, 39 of the manually modified (Group 1 and 2) and 21 of the newly designed SPs (Group 3) were successfully cloned and tested for amylase activity in the MTP assay, showing substantial difference compared to their original counterparts (Groups 1 and 2) and very effective secretion (Group 3), respectively (Figure 4b). Remarkably, out of the 21 in silico pseudo-randomly designed SPs (Group 3), 5 showed a secretion efficiency higher than AmyQ with its native SP.

To further validate the quality of the model in predicting SP efficiency for AmyQ secretion (i.e. WA value), we selected 5 SPs from each of the three groups and analysed their behaviour in the NLR-based assay. Dot plots of two variants with the respective original (‘ORIG’) and manually edited (‘EDIT’) versions are shown in Figures 4c and 4d, respectively, and highlight the clear shift in secretion efficiency for the two versions depicted in Figure 4a and 4b. Figure 4e summarizes the WAs of the 15 selected SPs and compares them with the WAs obtained at each step of the workflow (i.e. library screening, model predictions and validation). Remarkably, 11 of the tested SPs fell within one unit of difference (i.e. +/-1 WA) from the predicted value, implying that the proposed workflow is indeed a powerful tool to quantify the efficiency of engineered SPs.

The amylase quantification experiments in MTP cultivations show that, although the model was trained only with NLR-based data, its predictions mostly retain their validity at larger scale. To our knowledge, we have thus achieved an unprecedented accuracy in the prediction of SP efficiency and present the first example of successful model-driven *de novo* design of highly effective SPs. In fact, *in silico* SP design based on our trained model already proved very effective in the present proof-of-principle study, since the best predicted SPs turned out to direct high-level secretion.

Altogether, we conclude that our approach can detect and explain the relevant SP features that influence the efficiency of protein secretion. It sets the stage for *in silico* tuning and *de novo* design of SPs. Although we limited our present study to one protein, the workflow can be extended to other industrially or biomedically relevant POIs by applying different enzymatic assays (56) and novel HT analytical systems (17, 57, 58). For the future, we advocate an iteration of our approach to obtain further insights into the general features that influence protein secretion. This may be achieved either by using a fractional factorial design (59, 60) to ameliorate the design space (e.g. combining regions with differently modified features rather than editing one at a time), or through the screening of different POIs (e.g. fused to the same set of SPs). Datasets thus obtained will further improve the generalizability and reliability on prediction and design of SPs directing high secretion levels. As a result, far smaller numbers of SPs will be screened, or SP sequences will directly be designed in silico, for instance with our pseudo-random approach, or by exploiting a novel ML-based tool for SP generation (61). We are therefore confident that, with less experimental testing, our approach will deliver a deeper understanding of SP function and more accurate, better tunable, and highly productive protein secretion systems.

## DATA AVAILABILITY

Supplementary Tables, Supplementary Figures, Supplementary Files, train and test data, the ML model, and the SHAP explainer have been deposited on Zenodo with the following doi: 10.5281/zenodo.6448806 and are available at https://doi.org/10.5281/zenodo.6448806. Additionally, an interactive Jupyter notebook to fully explore the model is available in the same repository.

Raw Illumina reads have been deposited on NCBI SRA under BioProject accession number: PRJNA825264 (data will be made publicly available upon publication; reviewers can access data at the following link: https://dataview.ncbi.nlm.nih.gov/object/PRJNA825264?reviewer=o1cud8f26678bl6jltegirr2vf.

Flow cytometry data have been deposited on FlowRepository under the experiment ID FR-FCM-Z57A at the following link: https://flowrepository.org/id/FR-FCM-Z57A

## ACKNOWLEDGMENTS

We would like to thank Nivitha Punniyamoorthy and Steven Schmitt for the technical support provided during the long hours spent at the Biosorter. We would also like to thank Irsan Kooi, Marcel Hillebrand, Rianne van der Hoek, and Ana Bulović for the fruitful discussions.

## AUTHOR CONTRIBUTIONS

J.M.vD., M.Hel., S.P., A.M., and T.v.R. conceived the project. S.G. and P.Z. designed the libraries. S.G. designed and performed cloning and transformation. V.D. and R.P. designed and optimized the NLR-based enzymatic assay. S.G. and V.D. performed the screening assay. S.G. prepared the sequencing library and analyzed the NGS data. S.G. and M.Hen. generated and explained the ML model. V.D. validated the assay and the model. S.G. and V.D. drafted the manuscript. S.P., J.Mv.D., A.M., and T.v.R. supervised the project and revised the manuscript.

